# Large-scale integrative taxonomy of the smallest insects reveals astonishing temperate diversity

**DOI:** 10.1101/2025.09.17.676869

**Authors:** Catherine Hébert, Colin Favret

## Abstract

Fairyflies (Hymenoptera: Chalcidoidea: Mymaridae) are a diverse but taxonomically neglected group of parasitoid wasps that attack eggs of other insects. Being among the very smallest of all insects, they are often overlooked in biodiversity surveys despite being one of the most abundant microhymenoptera in many habitats. The traditional approach of morphological sorting for species delimitation can be challenging due to their minute size and meticulous slide-mounting technique. Ways to accelerate their discovery are needed. We conducted the first large-scale study of Mymaridae in temperate forests, combining DNA megabarcoding and the Large-scale Integrative Taxonomy (LIT) workflow to describe their diversity. We obtained COI barcodes from 2,098 specimens and used ASAP and RESL for species delimitation. Between 42 and 114 molecular clusters were delimited. Reducing morphological validation to only 9% of the sample enabled accurate determination while limiting time and effort. We confirmed the presence of 55 species, including many potentially new to science. The LIT workflow was effective for Mymaridae, although cryptic diversity remains unresolved in some large clusters, especially in the genera *Alaptus* and *Anagrus*, where high haplotype diversity and morphological ambiguity suggest additional hidden species. DNA reference databases proved unreliable, with less than 1% correct species matches, highlighting the taxonomic gap for this group. Nonetheless, we contributed 16 new identified reference barcodes to public databases and added new provincial and national species records for Canada. Our results demonstrate the value of combining molecular and morphological data in a standardized workflow and underscore the importance of improving reference databases for effective biodiversity assessments of dark taxa like microhymenoptera.

## Introduction

Hyperdiverse taxa such as insects are poorly known, especially within Hymenoptera and Diptera (Srivathsan et al. 2023). Moreover, Hymenoptera is estimated to be the most diverse insect order, mainly due to the immense richness of parasitoids (Forbes et al. 2018). Parasitoid wasps play a critical role in regulating the populations of their hosts, many of which are pests in agriculture and forestry. These wasps can also serve as valuable biodiversity indicators within arthropod communities (Anderson et al. 2011). Among them, the Mymaridae Haliday, also known as fairyflies, are of particular interest due to their ecological importance as egg parasitoids, mainly of Hemiptera, Coleoptera, and Psocoptera (Huber 1986). With over 1,600 named species worldwide (Nastasi et al. 2024), these smallest of insects are the sister group to the rest of the Chalcidoidea (Cruaud et al. 2023). Despite their abundance in many habitats (Vieira et al. 2013, Dall’Oglio et al. 2016, Haas-Renninger et al. 2023, 2024, Wang et al. 2024), and especially in forest environments (Vance et al. 2007, Quicke et al. 2023), mymarids remain overlooked in biodiversity surveys, primarily due to their minute size, lack of distinctive coloration, and other taxonomic challenges (Huber 1986). Many fairyfly species are difficult to identify based on morphology alone, and the labor-intensive slide-mounting process required for species-level identification further complicates large-scale studies. Reliable taxonomy is necessary for parasitoid wasp conservation and for their use in biological control (Bickford et al. 2007, Driesche and Hoddle 2016), but this “dark taxon” (Page 2016, Hausmann et al. 2020), with substantial gaps in knowledge regarding its diversity, distribution, ecology, and host preferences, is a victim of the taxonomic impediment (Ebach et al. 2011, Engel et al. 2021).

Traditional taxonomic workflows in the study of insects rely on morphospecies identification followed by DNA barcoding (Hebert et al. 2003), a process that can be biased by misidentifications at the level of the morphological OTU (Operational Taxonomic Unit) (Krell 2004, Derraik et al. 2010). This approach is particularly problematic in the case of cryptic or overlooked species such as mymarids, where morphological traits can be difficult to discern, and where sexual dimorphism can lead to male and female specimens being separated into different OTUs. Indeed, cryptic species are common within Hymenoptera (Li and Wiens 2023). To overcome these issues, DNA barcoding has emerged as a powerful tool for species identification (Hebert et al. 2003), but challenges persist in generating comprehensive reference libraries. Current databases are sparse, with Mymaridae largely underrepresented in DNA reference collections such as the Barcode of Life Data Systems (BOLD; Ratnasingham and Hebert 2007), where only 24 out of the 94 named species recorded in Canada are represented (Huber et al. 2021). Furthermore, many undescribed species exist, as evidenced by the number of Barcode Index Numbers (BINs) in BOLD relative to the known species diversity (Bennett et al. 2019). This lack of representation can impact biodiversity estimates obtained from studies employing metabarcoding or eDNA (Remmel et al. 2024, Kilian et al. 2025) which are already affected by the small size of certain insects such as fairyflies. There is thus a need to add well-identified and well-curated mymarid DNA barcodes to reference databases.

DNA barcoding, primarily developed for species identification, is now commonly used to group specimens into putative species or molecular OTUs (MOTUs), also known as molecular species delimitation (Blaxter 2004). Many algorithms have been developed for this purpose, either based on genetic distances or phylogeny. Distance-based methods can use an a priori genetic threshold to delimit clusters, like Objective Clustering (OC; Meier et al. 2006) and Refined Single Linkage (RESL; Ratnasingham and Hebert 2013), or they can infer a barcoding gap from the data itself, like Assemble Species by Automatic Partitioning (ASAP; Puillandre et al. 2021). Tree-based methods such as General Mixed Yule Coalescent (GMYC; Pons et al. 2006) and Poisson Tree Processes (PTP; Zhang et al. 2013) model speciation using divergence times or nucleotide substitution rates. Even if DNA-based methods could alleviate the drawbacks of morphological sorting, an approach combining multiple sources of data for species delimitation, also known as integrative taxonomy (Dayrat 2005, Padial et al. 2010), is still preferred.

Recent advances in high-throughput DNA barcoding, or megabarcoding (*sensu* Chua et al. 2023), and the development of a reverse workflow for species delimitation offer new opportunities (Puillandre et al. 2012, Kekkonen and Hebert 2014, Meier et al. 2016, Wang et al. 2018, Hartop et al. 2022). A reverse workflow removes prior morphological sorting, instead relying on molecular data to hypothesize species boundaries, which are then afterwards tested using morphological traits. This integrative approach, exemplified by the Large-scale Integrative Taxonomy (LIT) protocol introduced by Hartop et al. (2022), proposes specific criteria to predict incongruence in molecular clustering and to select which specimens need further evaluation, thereby accelerating the species hypothesis validation process. The studies that have taken advantage of molecular data in species delimitation within Mymaridae address only small species groups or species complexes (Triapitsyn, Rugman-Jones, et al. 2010, Zanolli et al. 2016, Nugnes et al. 2017, Triapitsyn et al. 2018, 2020, 2021, Triapitsyn, Kusuhara, et al. 2023, Triapitsyn, Rugman-Jones, et al. 2023). Little is known about the effectiveness of current DNA-based species delimitation methods for larger and more diverse samples of these dark taxa. The LIT approach has proven successful in the study of certain insect groups, especially Diptera (Hartop et al. 2022, Caruso et al. 2024, Riccardi and Hartop 2024, Meier et al. 2025), but has yet to be applied to other challenging taxa, such as parasitic hymenopterans.

In this study, we employ DNA megabarcoding, combined with molecular species delimitation and morphological analysis, to investigate and describe the diversity of Mymaridae in Quebec’s temperate forests. By sequencing over 2000 specimens and using LIT criteria, we built a DNA barcode reference library and tested the efficacy of two commonly used distance-based algorithms, ASAP (Puillandre et al. 2021) and RESL (Ratnasingham and Hebert 2013), in delimiting mymarid species. We conclude that the reverse workflow and LIT is a viable approach for elucidating the diversity of mymarids and parasitoid Hymenoptera in general.

## Materials and methods

### Field sampling

We collected insects using Voegtlin-style suction traps (Fig. 1A; Favret et al. 2019). These traps are particularly useful for capturing a large diversity and abundance of very small hymenopterans. Insects were collected in a mason jar containing propylene glycol (commercial antifreeze). Antifreeze evaporates less than alcohol while preserving DNA and specimen morphology (Moreau et al. 2013, Martoni et al. 2021). The traps ran for a week at a time. Then the samples were gathered, and the traps removed or restarted with a fresh battery. Each mason jar contained a label with a unique identifier, the sampling site number, and dates.

**Figure 1.**
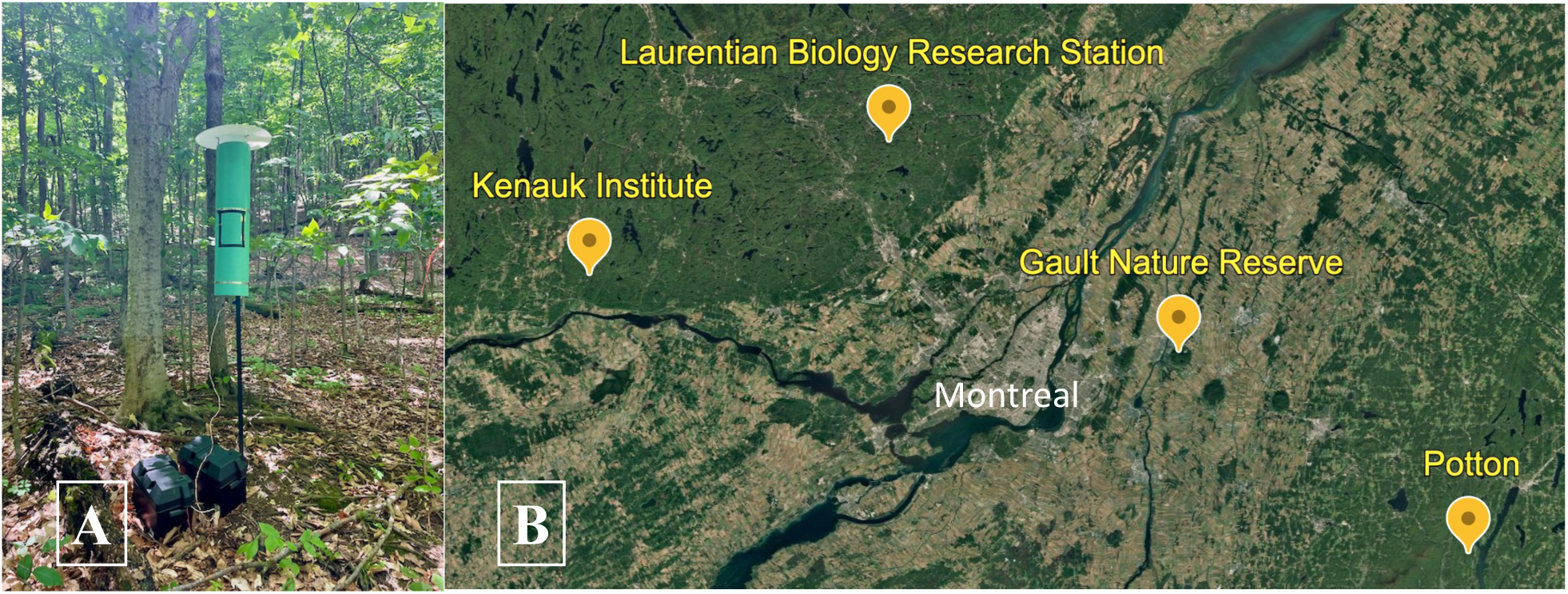
A) Example of a Voegtlin-style suction trap installed at the Gault Nature Reserve B) Collection sites of the four forested environments in Quebec. We collected a total of 70 week-long samples that were sorted first to the ordinal level, then parasitoid wasps were further identified to the family level. The identifications were done using identification keys from Goulet and Huber (1993), Gibson et al. (1997), Noyes (2019), and the wasp classification book from Nastasi et al. (2024). Specimens were transferred from propylene glycol to 95% ethanol and kept at -20°C to preserve DNA. Samples from the weeks of 2023-06-23 to 2023-08-15, which include the sampling of all ten sites, were selected for this study. These specimens are vouchered, databased, and deposited in the Université de Montréal’s Ouellet-Robert entomological collection (QMOR). As in Hébert et al. (2022, 2023), specimen data in Darwin Core format are also published through Canadensys and GBIF to facilitate its accessibility (Hébert and Favret 2025).

We sampled ten sites distributed in four administrative regions of Quebec (Fig. 1B), dominated by mixed-deciduous or sugar maple forests (Table 1). Sites 1 and 2 (Table 1), located at the University of Montreal’s Laurentian Biology Research Station (SBL), were previously sampled in 2015 and 2021 for other projects (Favret et al. 2019, Kalboussi and Favret 2022). Traps at these two sites were run for the whole 2023 growing season, from May through October. The other eight sites were sampled for two or three weeks each from mid-June to mid-August, the peak of mymarid abundance and diversity (unpublished data). The traps were at least 50 meters apart. Authorizations were acquired before sampling.

**Table 1.**
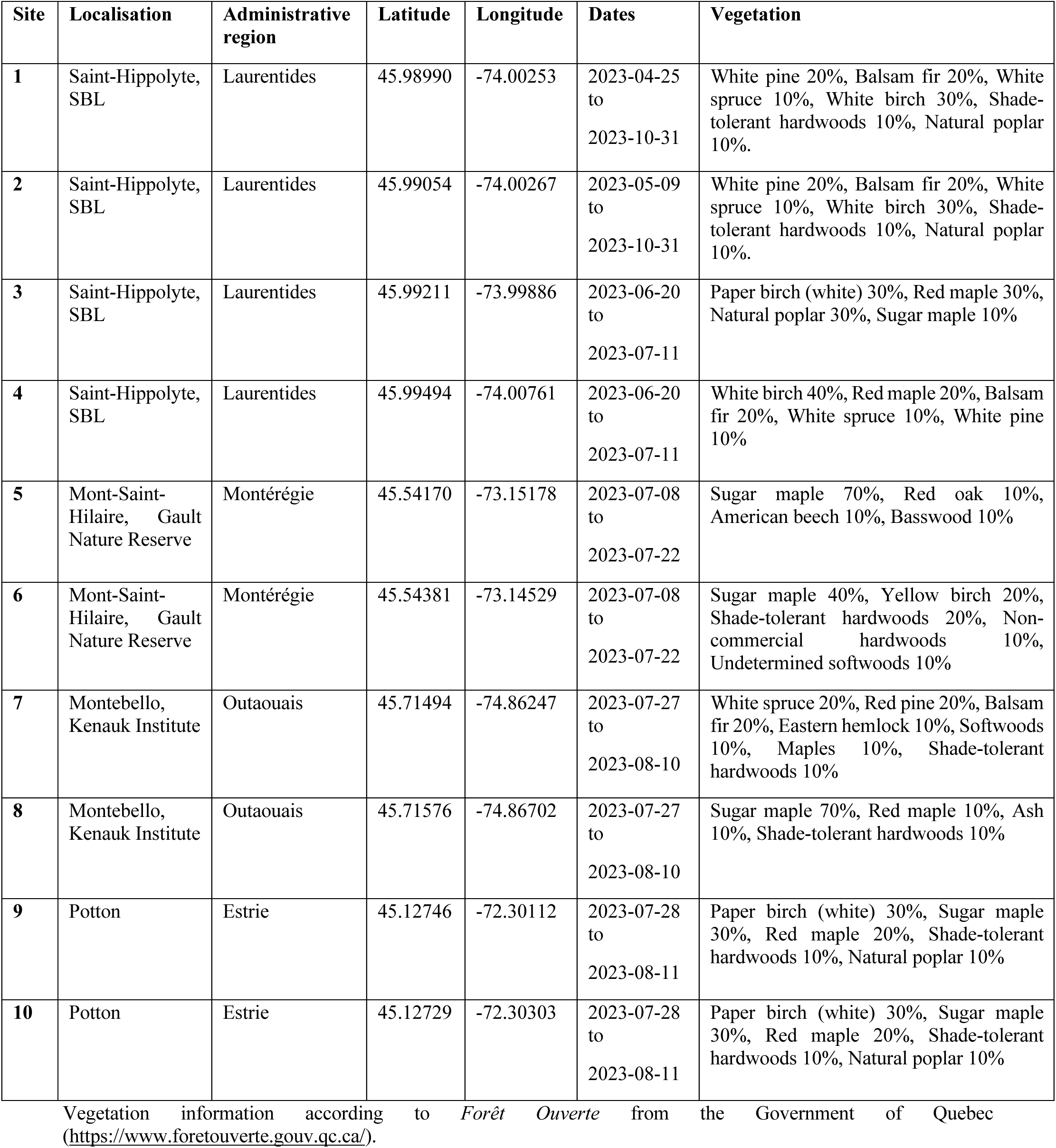
Collection sites in Quebec, Canada, collection dates and vegetation characteristics.

### High-throughput DNA sequencing

#### DNA extraction

Genomic DNA was extracted non-destructively from 2,138 individual mymarid specimens with the QuickExtract™ DNA Extraction Solution (Lucigen, New York, USA). First, each specimen was put in a 0.2 mL tube left open until the alcohol evaporated. Fifteen µL of QuickExtract solution was then added to each tube which was kept overnight in a thermocycler at 65°C, followed by a 98°C enzyme denaturation treatment for two minutes. The incubation time was much longer than what is recommended by the manufacturer, but other research (Martoni et al. 2021) and preliminary tests confirmed that the DNA yield was higher. The DNA extracts were pipetted into 96-well microplates and kept at -20°C. The tubes containing the specimens were filled with 70% ethanol and kept at -20°C for later morphological analyses. DNA quantification and absorbance ratio were measured for a subset of each plate using a spectrophotometer (NanoDrop Technologies, Wilmington, USA).

#### PCR amplification and Illumina sequencing

PCR amplifications targeted a 316-bp region of the cytochrome c oxidase 1 (COI) gene, covering the 3’ end of the “Folmer region”. Such mini barcodes are easier to amplify and work as well as longer barcodes for species delimitation (Yeo et al. 2020). We used the new primer combination BF1 (5’-ACW GGW TGR ACW GTN TAY CC-3’) (Elbrecht and Leese 2017) and C_LepFolR (5’-TAA ACT TCW GGR TGW CCA AAA AAT CA-3’) (Elbrecht et al. 2019). Five-bp tags, designed by Glenn et al. (2019) and containing one- to four-base heterogeneity spacers, were added to the primers for multiplex sequencing. The spacers help in the sequencing of low complexity libraries (Fadrosh et al. 2014). Illumina sequencing adapters were also attached to the primer tags resulting in fusion primers ranging from 59 to 69 bases. A combinatorial indexing approach was used with the generation of 12 forward and 8 reverse primers to make 96 unique combinations.

PCR reactions were carried out in 96-well plates, each well containing 10 µL of Phire Green Hot Start II PCR Master Mix (Thermo Scientific, Waltham, MA), 2 µL of 5 nM of each fusion primer, 1 µL of DNA template and water up to 20 µL. A gradient PCR protocol was first used to determine the optimal annealing temperature for the primer combination. Final PCR conditions were an initial denaturation at 95°C for 3 min, followed by 35 cycles of denaturation at 95°C for 30 sec, annealing at 47°C for 30 sec, and extension at 72°C for 2.5 min, and a final extension at 72°C for 5 min. A subsample of ∼12 amplicons per plate was checked by electrophoresis on a 1% agarose gel containing GelRed (Biotium, Fermont, USA). Two µL of each of the 96 amplicons from a plate were pooled and 20 µL of this solution was sent to the *Centre d’expertise et de services Génome Québec* (CESGQ) for final library preparation and sequencing of 2×300 paired end reads on an Illumina NextSeq2000 instrument. Since each of our pools possessed the same tagging combinations, the CESGQ added a second round of unique indexes to allow for the subsequent demultiplexing of libraries.

A second round of PCR was done for the 74 samples that failed to produce a DNA sequence at the end of the process. The same conditions were used except that 2 µL of DNA template instead of 1 µL. In total, 24 libraries containing between 51 and 96 DNA samples each were sent for Illumina sequencing.

#### Bioinformatic pipeline

Read quality was assessed with the FASTQC program available freely on the Galaxy platform (usegalaxy.org). The following analyses were done with Geneious Prime (version 2023.2.1). FASTQ files from both read directions were imported and set to paired reads. Demultiplexing was done with the function “Separate reads by barcode” for both forward and reverse tags, allowing no mismatches. Primer sequences were removed with the “Trim Ends” function allowing for five mismatches, a minimum overlap of five and an error probability limit of 5%. The BBDuk (version 38.84) trimmer was used to trim Illumina adapters, ends of low quality (<20Q) and sequences shorter than 250-bp. The reads were then merged using BBMerge (Bushnell et al. 2017). Demultiplexed reads were clustered at 98% similarity and minimum overlap of 250-bp with a “de novo assembly”. Several contigs were obtained and the consensus sequence of each one was extracted. Some sequences retained part of the primers and were manually edited when necessary. The consensus sequences were submitted to GenBank BLAST searches from Geneious and the top 5 hits displayed. Sequences not corresponding to Mymaridae were discarded. For some specimens, two or more consensus sequences belonged to Mymaridae. The contig with the maximum reads was selected. The others were considered possible cases of heteroplasmy or DNA cross-contamination and were discarded. We chose a read coverage threshold of 10x to retain the barcode. Consensus sequences were translated and examined for stop codons or frameshift mutations indicating possible pseudogenes, eight of which were then discarded.

All sequences generated can be found in the *Mymaridae of temperate forests of Quebec* dataset on BOLD (DS-MYMAQC; BOLD 2025) and on GenBank (accession numbers PX064719-PX066816). BOLD does not create new BINs based on barcodes less than 500 bases, but ours were nonetheless assigned to existing BINs when appropriate.

### Molecular species delimitation

Mymarid specimens were first clustered into putative species using molecular data. Two distance-based species delimitation approaches were selected and compared: Assemble Species by Automatic Partitioning (ASAP; Puillandre et al. 2021) and Refined Single Linkage (RESL; Ratnasingham and Hebert 2013). ASAP follows a hierarchical clustering approach to propose species partitions using pairwise genetic distances (Puillandre et al. 2021). Partitions are scored based on the probability of panmixia and the width of the barcoding gap; lower scores are usually better. ASAP was downloaded from the iTaxoTools bioinformatic platform (Vences et al. 2021). A multiple sequence alignment with MAFFT (Katoh and Standley 2013) was generated in Geneious Prime and used as input. Default parameters were selected in ASAP with the Jukes-Cantor distance. Three partitions with equally lower scores were kept for comparison.

RESL is the algorithm used by BOLD for their BIN system (Ratnasingham and Hebert 2013). It uses a single linkage clustering at a distance of 2.2% to generate first partitions, then refines them using Markov clustering. It was performed through the *Cluster sequences* tool from the BOLD workbench, with default parameters.

#### Cluster validation and identification

Validation of clusters followed the LIT protocol (Hartop et al. 2022). The guidelines proposed have never been used with Hymenoptera and thus were tested here to validate their utility. The partition with the smallest number of putative species was used as a baseline to compare with the other partitions. Clusters were categorized as potentially incongruent (PI) if they were either unstable, meaning that their composition differed between partitions, or had a high intraspecific genetic p-distance (>1.5%). For clusters not flagged as potentially incongruent, specimens representing the most divergent haplotypes were selected for morphological identification. Potentially incongruent clusters were checked more thoroughly, with the addition of specimens representing the dominant haplotype (containing >20% of the specimens). Even if most identification keys for Mymaridae address female specimens, both males and females were selected to match descriptions. When possible, at least two females of each were selected to mount in different views. If haplotypes matched only male specimens, the closest one that had females was also chosen to permit the identification. If multiple species were found in one cluster, it was split to match another partition and additional specimens representing these new clusters were chosen for validation. If no partitions matched morphology, a conservative approach was adopted, and the clusters were left as unresolved.

To better visualize clusters, minimum spanning haplotype networks were generated using Hapsolutely (Vences et al. 2024) downloaded from the iTaxoTools platform. TaxI2.2 (Steinke et al. 2005) was used to calculate intra-*vs*. inter-specific distances based on uncorrected p-distances.

Specimens selected for mounting were first photographed with a Keyence VHX-7000N Digital Microscope. They were then slide-mounted in Canada balsam following Huber (2015) for morphological identification, with one minor change to the protocol: terpineol was used instead of clove oil.

Morphological determinations were made using various identification keys that are listed in Hébert and Favret (2025). Keys outside of the North American range as well as original descriptions were also used. Determinations were compared with identified specimens available at the Canadian National Collection of Insects, Arachnids, and Nematodes (CNC) in Ottawa, Canada.

We acknowledge the use of ChatGPT-4 in the proofreading of this manuscript. All intellectual contributions and conceptual work presented herein are exclusively those of the authors.

## Results

### Sequencing results

Overall, we collected 8,907 parasitoid wasps representing 22 different families. Of those, 3,464 (39%) were Mymaridae. We extracted DNA from 2,139 mymarid specimens, and 2,113 sequences were obtained, for a sequencing success rate of 98%. Several sequences belonged to *Megaphragma* Timberlake (Chalcidoidea: Trichogrammatidae), mistakenly identified as Mymaridae in our original sorting, and some specimens yielded apparently contaminated sequences. We removed those 15 sequences, leaving a final dataset of 2,098 DNA barcodes, with 309 unique haplotypes.

### Large-scale Integrative Taxonomy

The three partitions with the lowest score from ASAP delimited 42, 83 and 93 mymarid clusters (Table 2), with an average of 4 haplotypes per cluster (range 1–50), and barcode gaps identified at 6.6%, 2.5% and 2.1%, respectively. These three delimitations are hereafter referred as ASAP42, ASAP83 and ASAP93. RESL found 114 clusters. With ASAP42 as a baseline, 20 clusters had a signature of incongruence between delimitation methods (PI clusters) (Table 3). Of those, five clusters were stable but had an intraspecific p-distance over 1.5%. The other 15 were both unstable and with high p-distances. Of the 22 clusters that were not flagged as PI, 13 were singletons. Mean intra-cluster genetic divergence was 1.37% (range 0−10.68%), and inter-cluster divergence was 15.43% (4.53−26.54%).

**Table 2.**
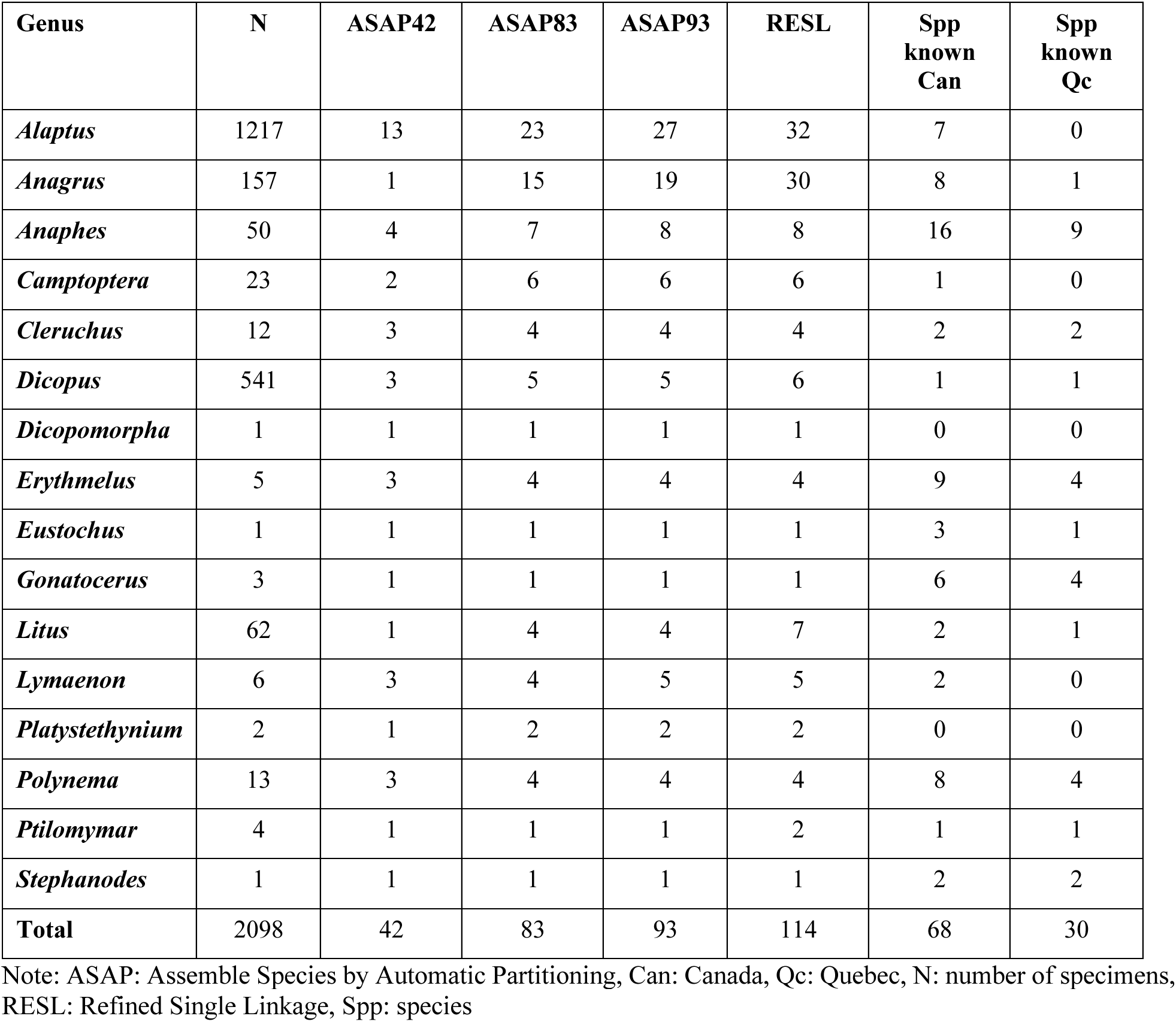
Abundance by Mymaridae genera, number of clusters for each delimitation method and species known for Canada and Quebec according to Huber et al. (2021).

**Table 3.**
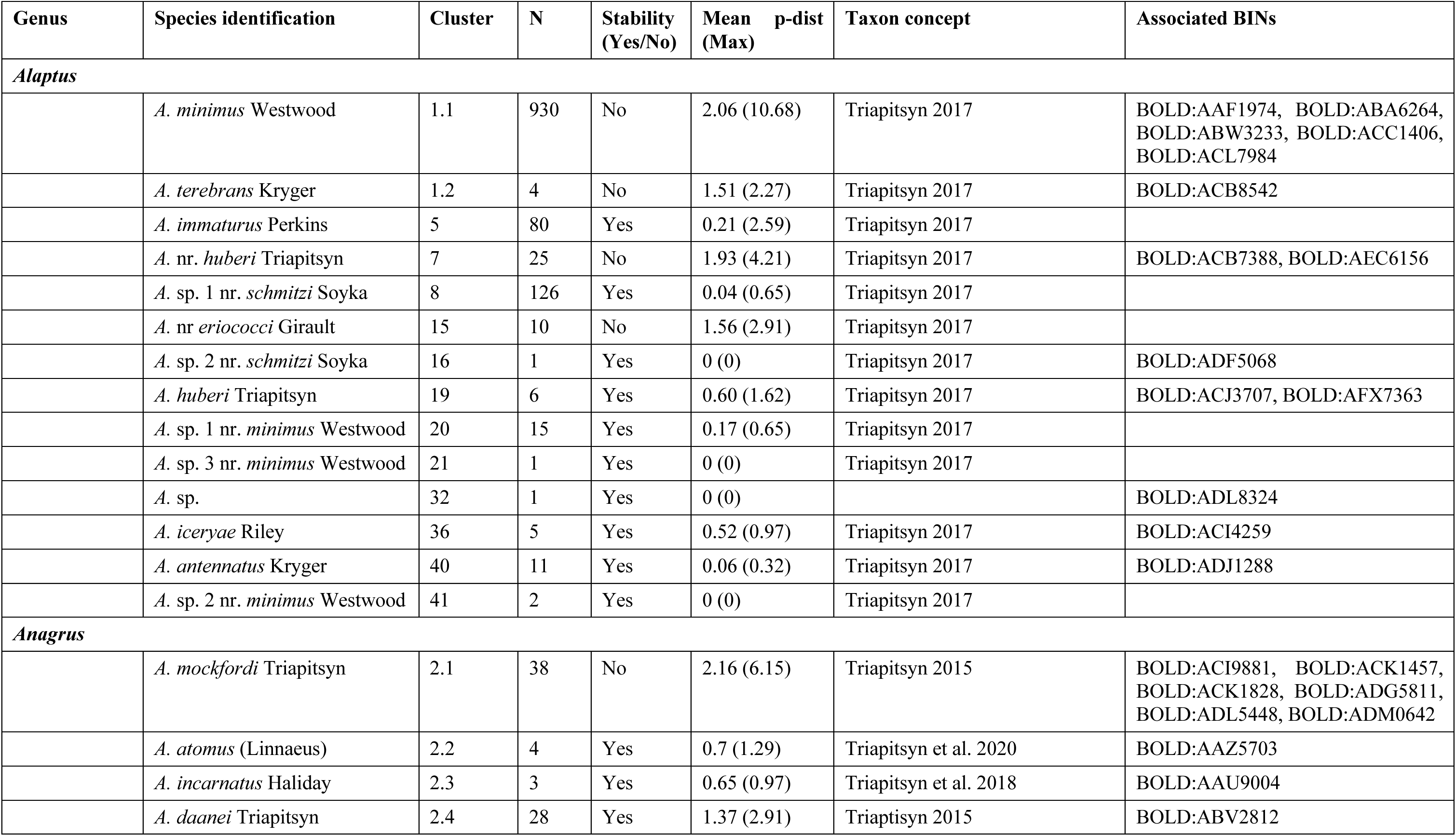
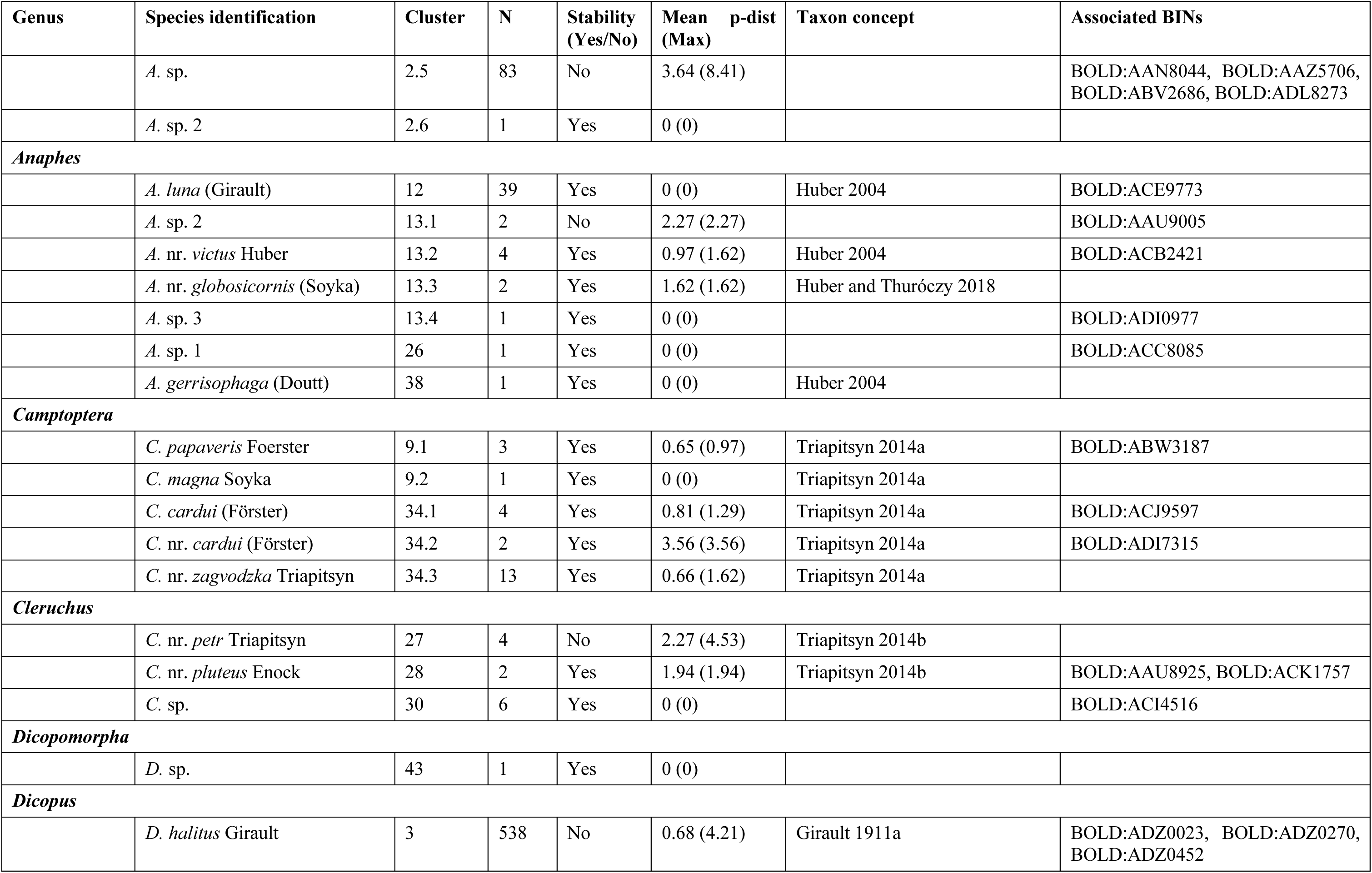
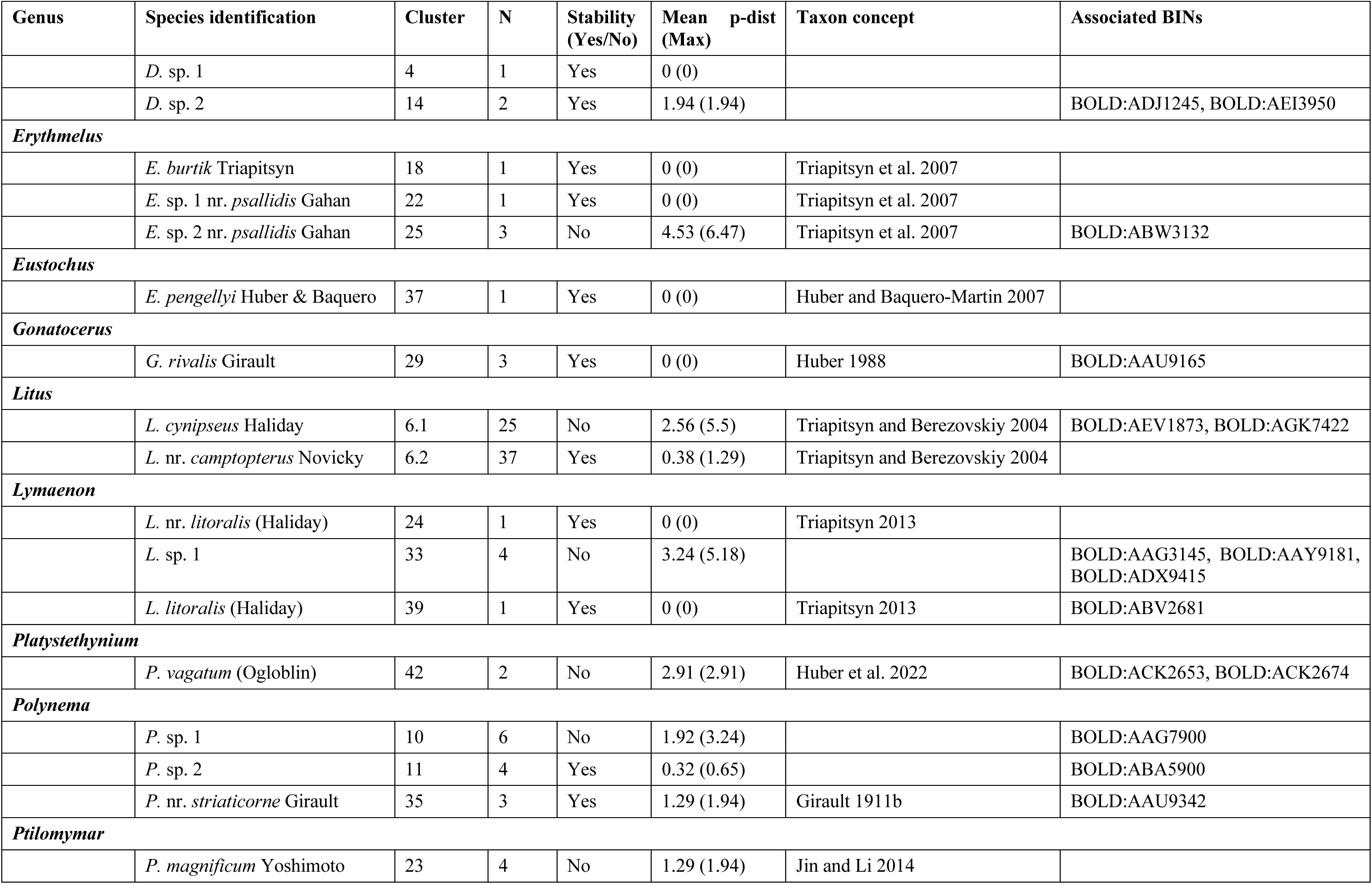
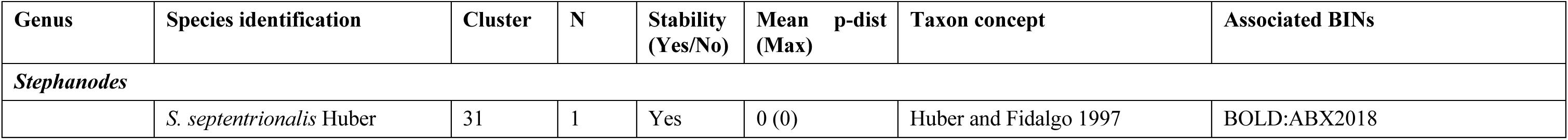
Species identification for the 55 delimited clusters of Mymaridae, LIT criteria of incongruence (stability and p-distance), taxon concept and associated BINs. QMOR specimen voucher data are available in Hébert and Favret (2025).

Of the 2,098 mymarid specimens, 185 individuals were mounted on slides for morphological validation, representing 9% of the total. We confirmed the presence of 55 mymarid species from 16 genera. Twenty-four clusters were confidently assigned to named species, 18 were morphologically close to named taxa but did not fully match their descriptions, and 13 could not be identified at all and might represent new species. Unflagged clusters, as well as stable ones with high divergence, were all found to be congruent with morphology. Of the rest of the PI clusters, nine were ultimately consistent with morphology, while six contained multiple species and were evaluated more thoroughly to determine if they matched another partition.

Genera *Cleruchus* Enock, *Dicopomorpha* Ogloblin, *Dicopus* Enock, *Eustochus* Haliday, *Gonatocerus* Nees, *Lymaenon* Walker, *Platystethynium* Ogloblin, *Polynema* Haliday, *Ptilomymar* Annecke & Doutt and *Stephanodes* Enock were well delimited by ASAP42. *Anaphes* Haliday and *Camptoptera* Foerster better matched the clustering of ASAP83. Unfortunately, cluster 10 from *Polynema* and cluster 33 from *Lymaenon* were both PI but contained mostly or only male specimens. They could not be confidently validated and were left as one cluster each. Genera *Alaptus* Westwood, *Anagrus* Haliday, *Erythmelus* Enock and *Litus* Haliday had more complex cases, as described below.

### Genus *Alaptus* Westwood, 1839

*Alaptus* was the most abundant genus in our sampling, representing more than 50% of the specimens (Table 3, Fig. 2). Morphological validation confirmed the molecular delimitation of 12 of the 13 clusters, including two that were flagged as PI. Cluster 16 was represented by only one male individual, but its nearest neighbor was 8.14% apart, thus confirming it was distinct even if we could not attribute it a species name. Cluster 1 was the only one of this genus that contained multiple species, with an intraspecific divergence of 11.33% prior to splitting. ASAP83 split cluster 1 into nine putative species but morphological validation was able to delimit only two groups: 1.1 and 1.2 (Fig. 2).

**Figure 2.**
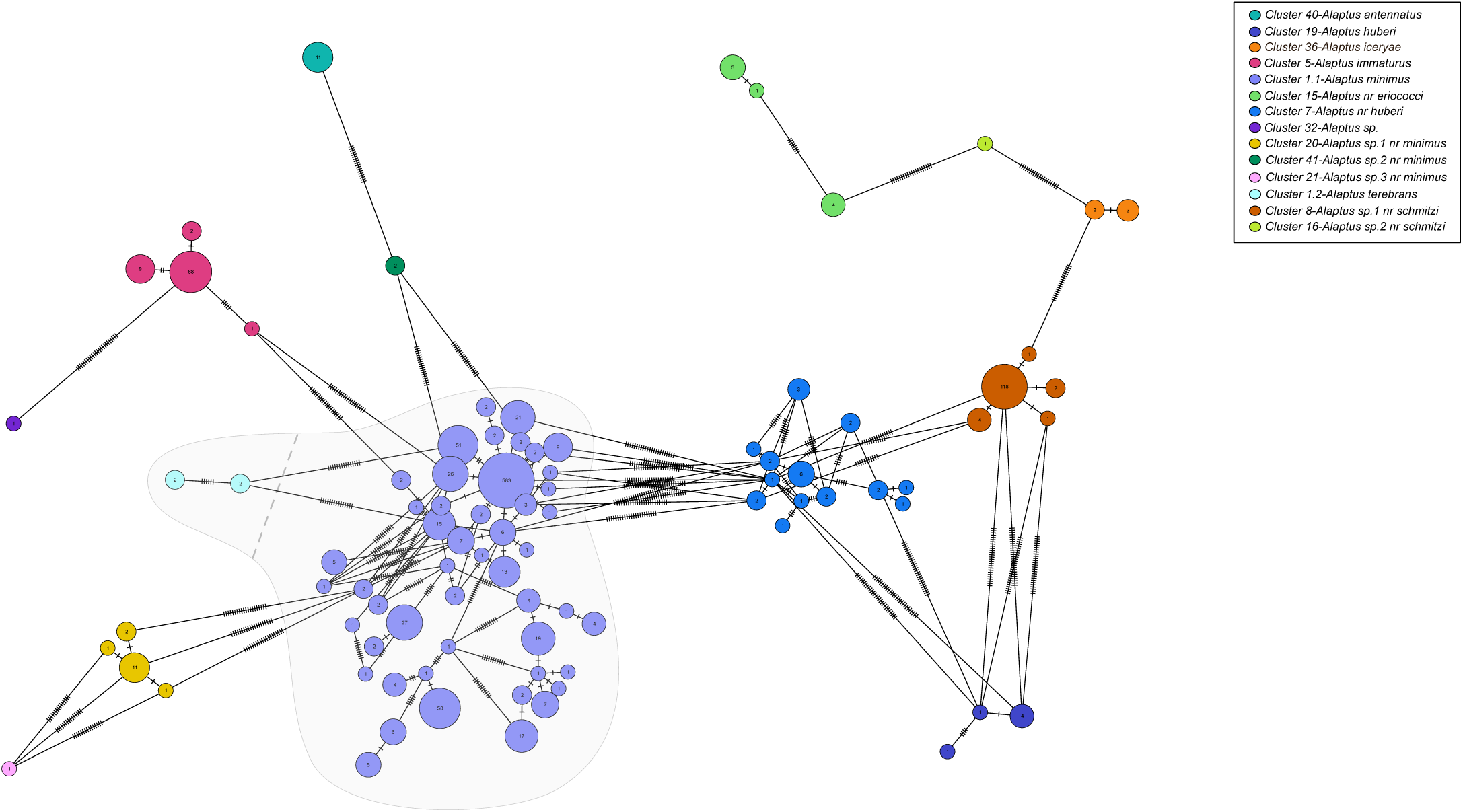
Haplotype network of *Alaptus* species. Nodes represent unique haplotypes and are color coded according to the species, number inside each node are the number of specimens possessing the haplotype in question, and the hash marks correspond to base pair differences between haplotypes. The grey background highlights clusters 1.1 and 1.2.

Most of the morphological variation observed in cluster 1.1 fell within the intraspecific variation recorded for the species *Alaptus minimus* Westwood (Fig. 3; Triapitsyn 2017). Some specimens had a higher ovipositor/metatibia ratio that corresponds better to *Alaptus fusculus* Walker, but every other important character matched those of *A. minimus*. Since we could not confidently separate them, these specimens were left in cluster 1.1 even if its intraspecific divergence was very high (up to 10.68%). Only cluster 1.2 was clearly separate from *A. minimus*, both genetically (mean distance of 7.22%) and morphologically, and was identified as *Alaptus terebrans* Kryger. It included only four male specimens but was associated with a BIN that had a female with a much longer ovipositor than *A. minimus* and *A. fusculus*. Our male individuals had several additional setae in the forewings that are diagnostic of *A. terebrans*, as well as fitting every other character described by Triapitsyn (2017), thus supporting its identification.

**Figure 3.**
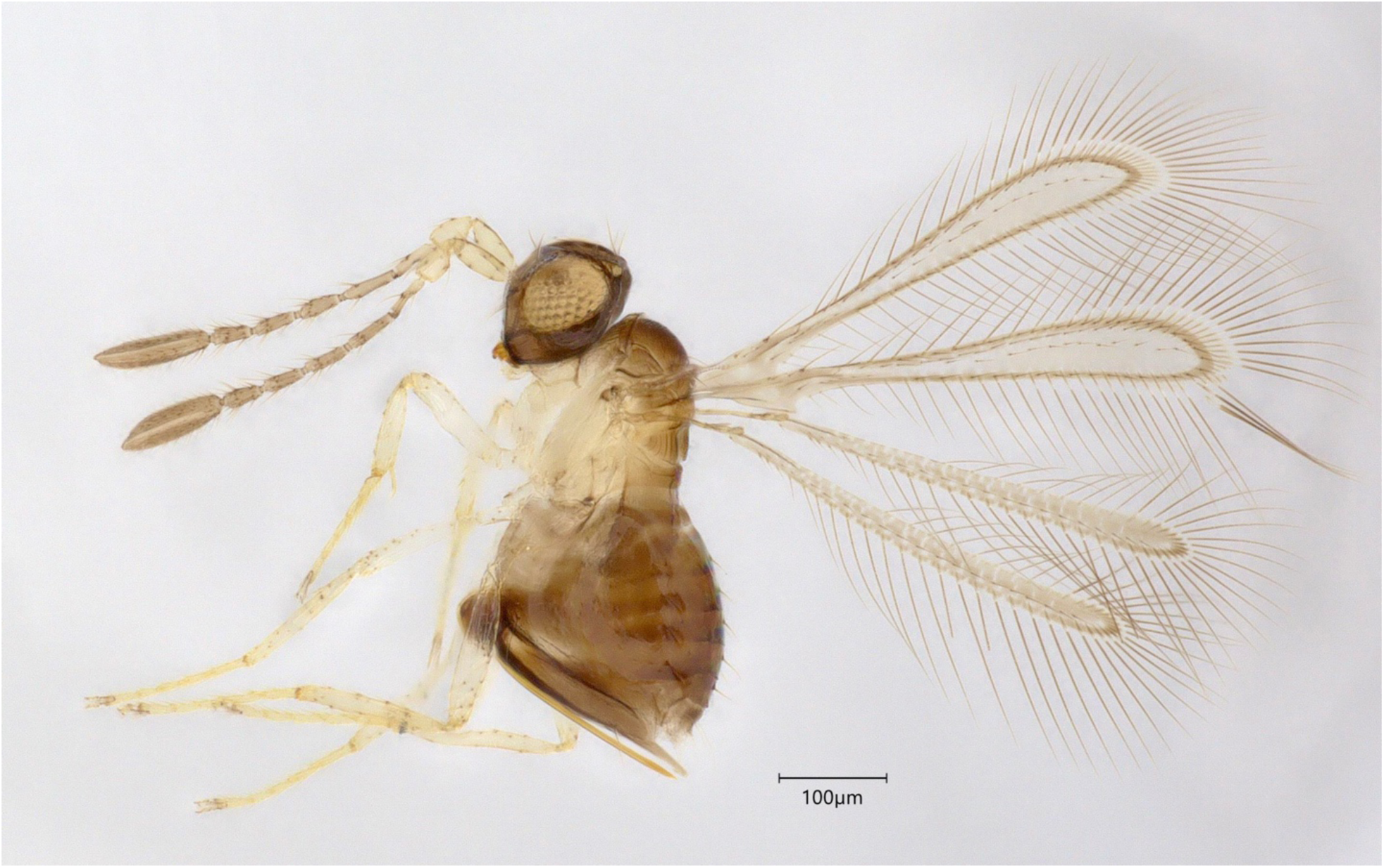
Representative of *Alaptus minimus* (QMOR84141).

Clusters 20, 21 and 41 all keyed to *Alaptus minimus* in Triapitsyn (2017) but had some ratios, mainly concerning antennal segments, that did not fit the intraspecific variation of this species. They all had clearly distinct barcodes (mean distances from 8.18% to 10.01%) from cluster 1.1 and were identified, morphologically, as species near *A. minimus*.

### Genus *Anagrus* Haliday, 1833

The 157 specimens of *Anagrus* were delimited as one putative species by ASAP42 that was correctly flagged as PI (Table 3, Fig. 4). However, morphology did not match the delimitation of ASAP83 which split it into 15 molecular clusters. Three clusters were identified as *Anagrus mockfordi* Triapitsyn (Fig. 5) and were grouped together despite their high intraspecific divergence (up to 6.15%). Four other clusters were well delimited with three of them corresponding to known species. The seven remaining clusters could not be confidently separated, even when looking at the ASAP93 partition. Many individuals, some with similar haplotypes, keyed nonetheless to different species in Triapitsyn (2015) but did not fully fit the descriptions. There was also overlap in some important characters between those specimens. Since no congruence could be found between molecular and morphological data, we decided to lump these seven clusters as one unidentified species.

**Figure 4.**
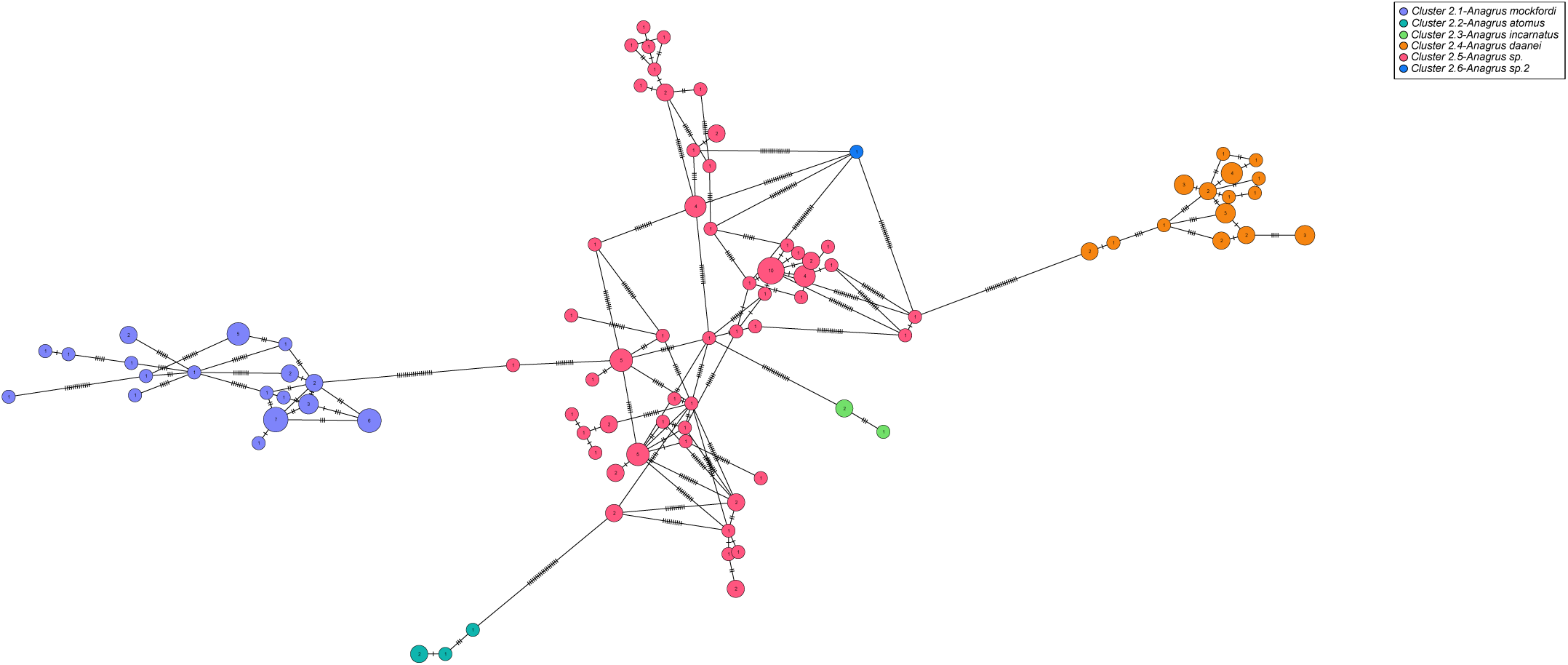
Haplotype network of cluster 2 with six species of *Anagrus*. Nodes represent unique haplotypes and are color coded according to the species, number inside each node are the number of specimens possessing the haplotype in question, and the hash marks correspond to base pair differences between haplotypes.

**Figure 5.**
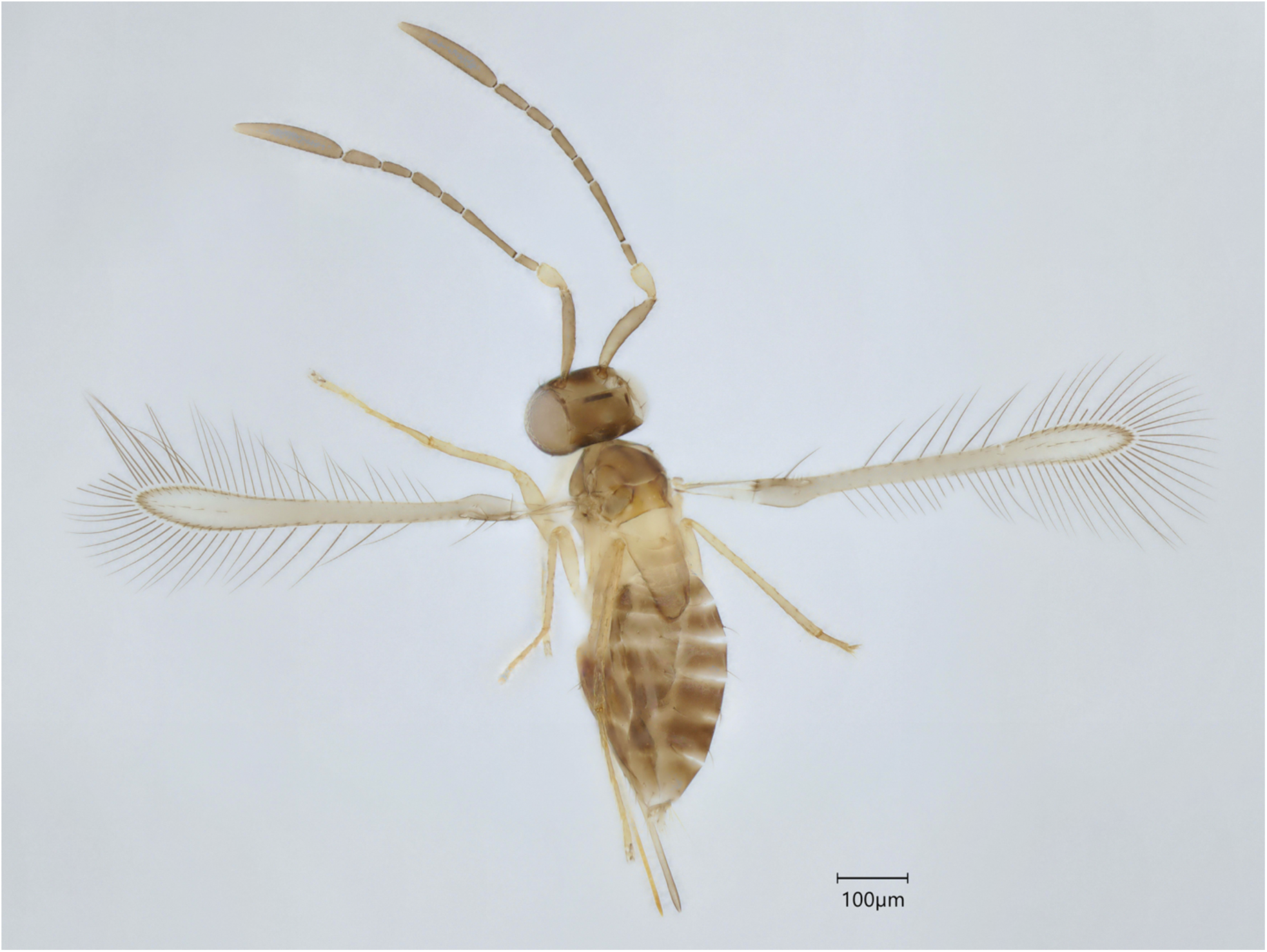
Specimen of *Anagrus mockfordi* (QMOR88129).

### Genus *Erythmelus* Enock, 1909

Cluster 18 was well defined as *Erythmelus burtik* Triapitsyn. Both clusters 22 and 25 (Table 3, Fig. 6) keyed to the species *Erythmelus psallidis* Gahan, with many of the characters measured overlapping between the two groups. However, they had a mean genetic distance of 10.57% from each other. Cluster 22 was composed of only one specimen, therefore morphological variation could not be evaluated. Cluster 25 was flagged as PI due to instability and high p-distance and one of the three specimens available had ratios regarding the ovipositor and the forewings larger than the description for *E. psallidis* (Triapitsyn et al. 2007) (Fig. 7). It also had more densely setose forewings than the other two individuals. Yet, this seemingly different individual was not the one that was split in a different cluster in ASAP83. Since this delimitation was not consistent with morphology, we decided to keep the ASAP42 clustering for this genus.

**Figure 6.**
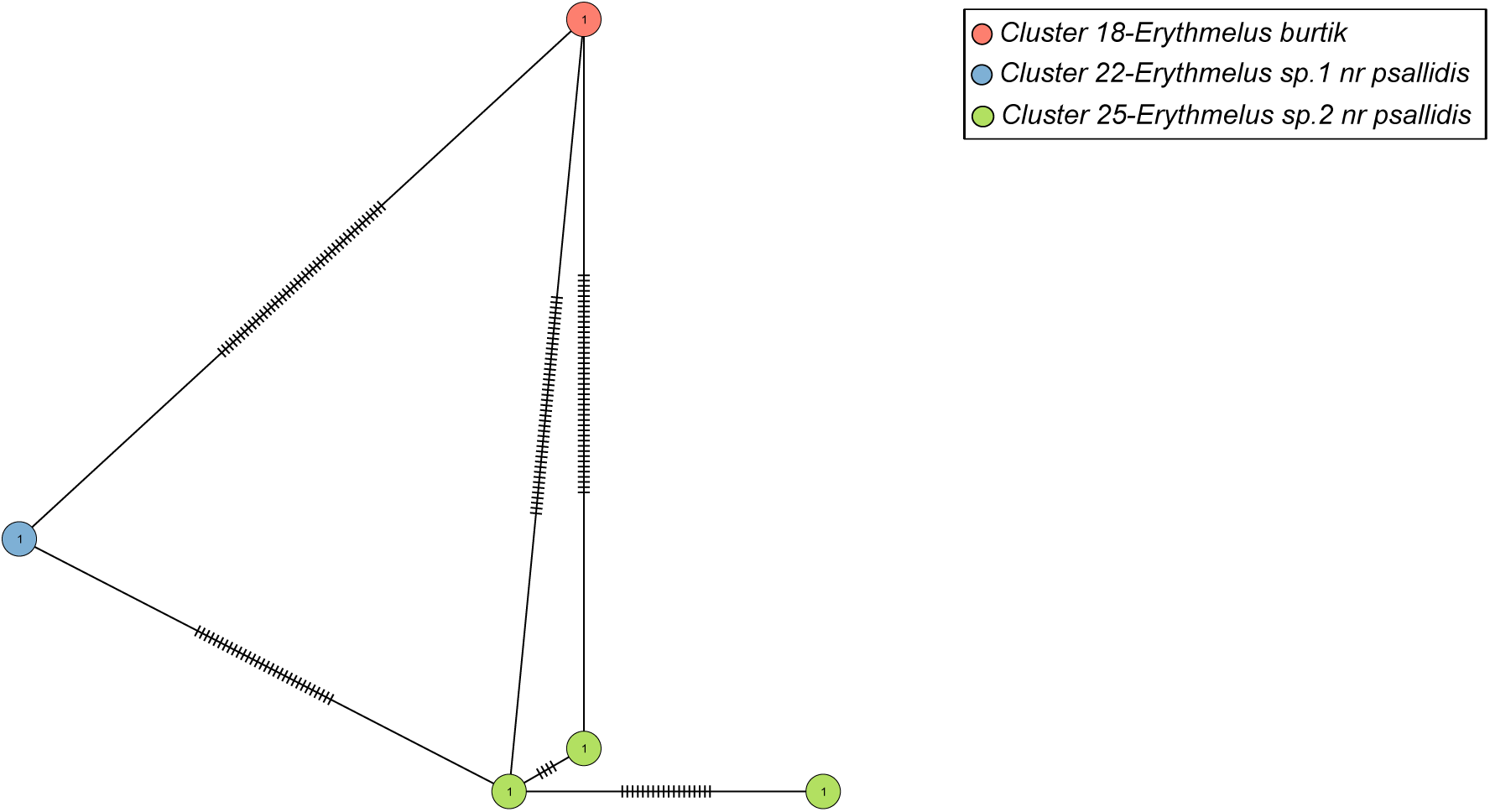
Haplotype network of *Erythmelus* species. Nodes represent unique haplotypes and are color coded according to the species, number inside each node are the number of specimens possessing the haplotype in question, and the hash marks correspond to base pair differences between haplotypes.

**Figure 7.**
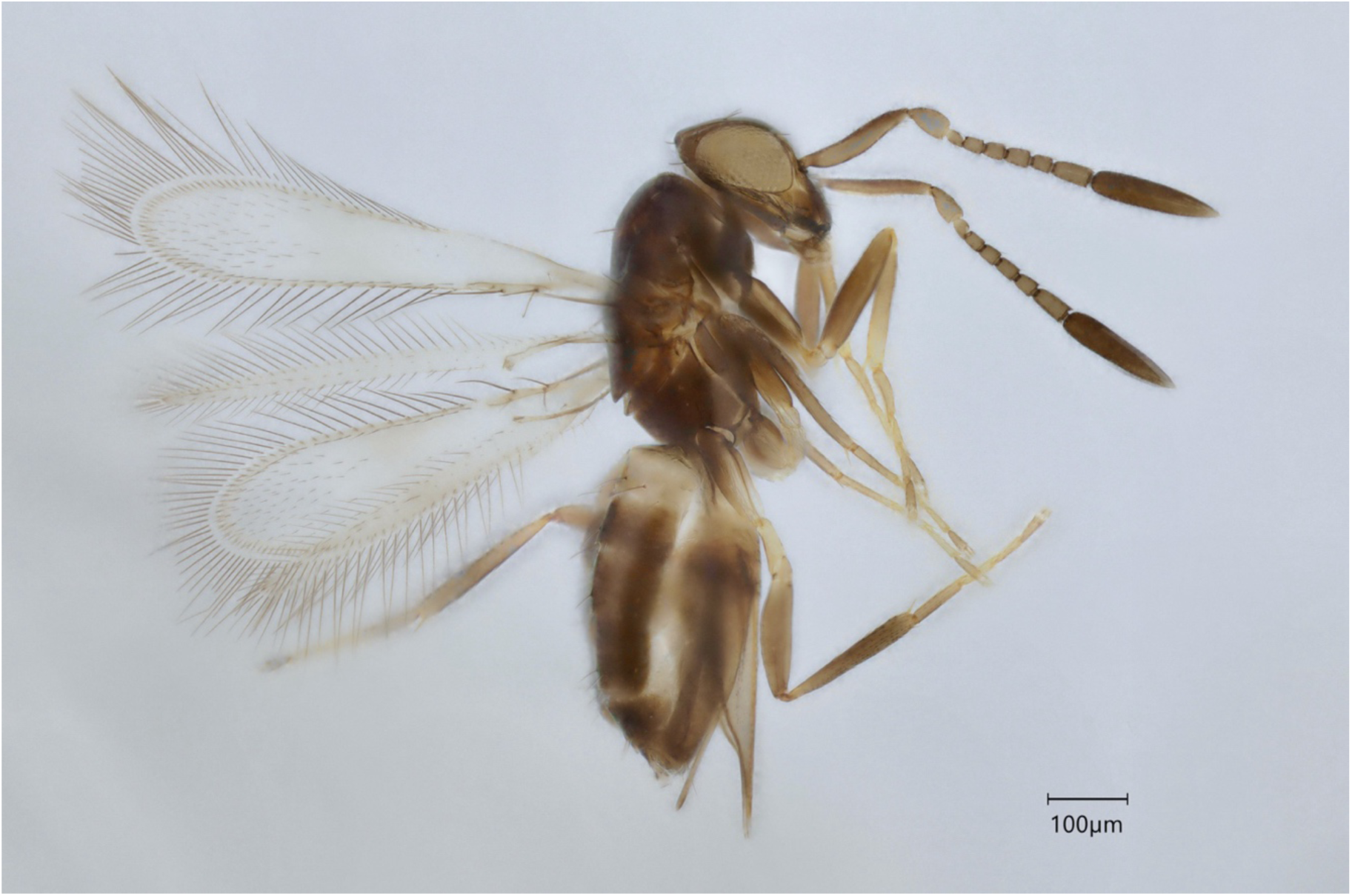
Representative of *Erythmelus* sp. 2 nr. *psallidis* (QMOR84176).

### Genus *Litus* Haliday, 1833

All *Litus* individuals were grouped as one PI cluster in ASAP42. Morphology confirmed the presence of multiple species (Table 3, Fig. 8) but could not validate the four clusters found by ASAP83. Cluster 6.1 contained three of the four groups from ASAP83 that were all identified as *Litus cynipseus* Haliday. Some variation was found in the pattern of setation of the forewings but there was an overlap in all other important characters such as the ratios regarding the antennal segments, the ovipositor and the forewings. Forewing chaetotaxy was thus considered intraspecific variation. Cluster 6.2 was identified as a species near *Litus camptopterus* Novicky (Fig. 9), which is known to occur in Quebec (Huber et al. 2021). Many of its diagnostic characters corresponded, except for the shorter length of the second funicular segment compared to the pedicel, which would incorrectly lead to *L. cynipseus* in Triapitsyn and Berezovskiy (2004).

**Figure 8.**
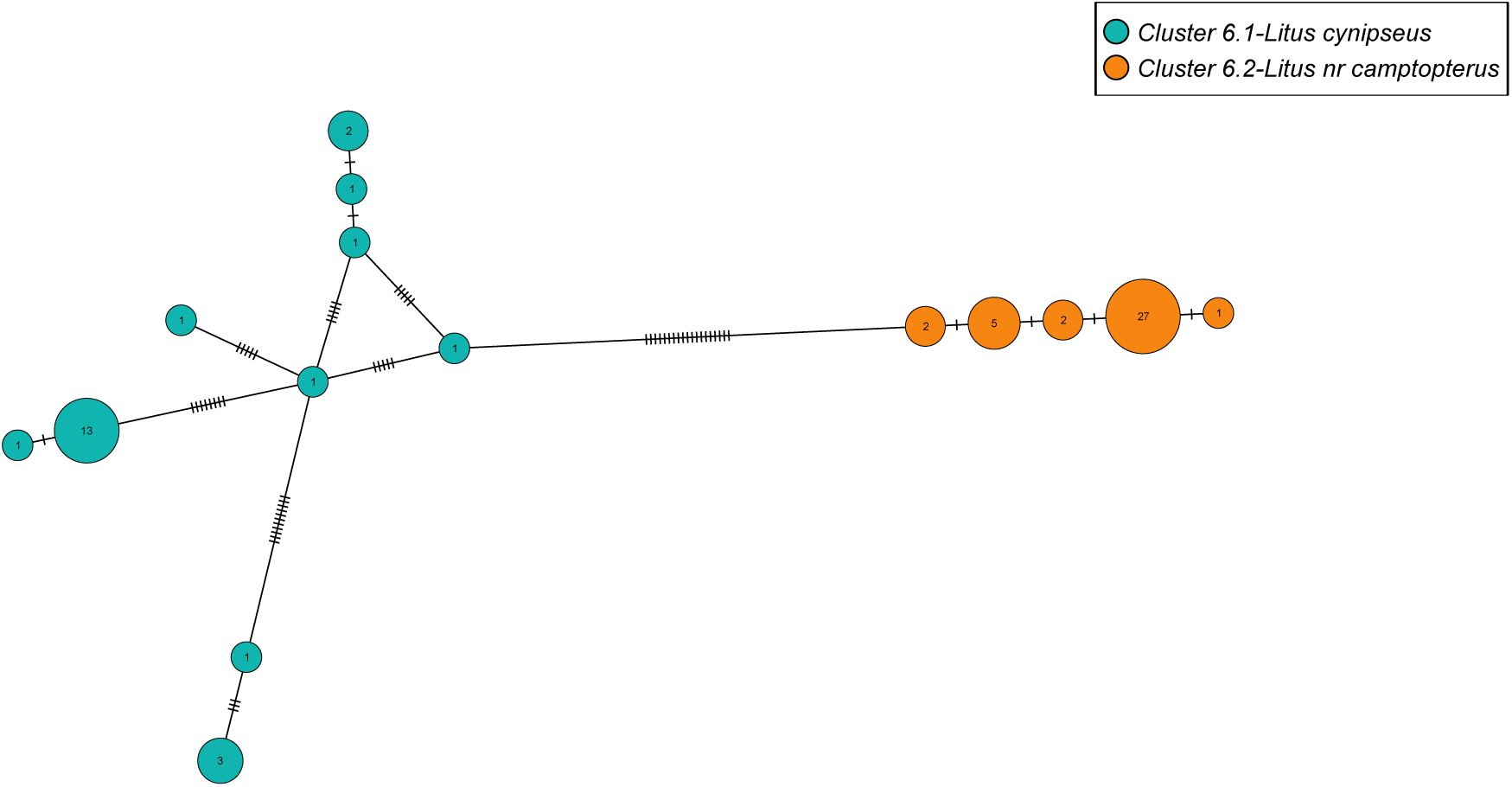
Haplotype network of *Litus* species. Nodes represent unique haplotypes and are color coded according to the species, number inside each node are the number of specimens possessing the haplotype in question, and the hash marks correspond to base pair differences between haplotypes.

**Figure 9.**
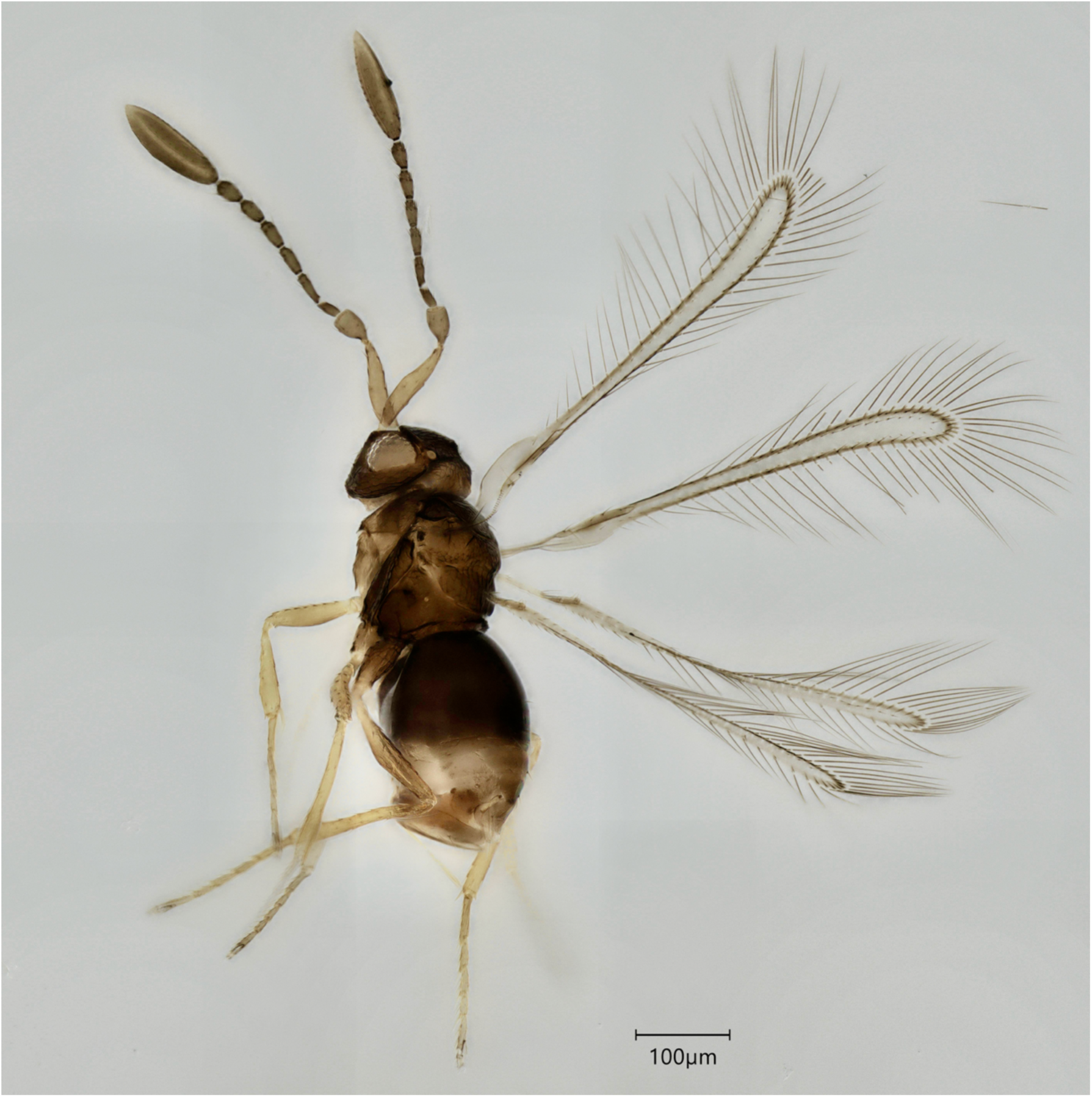
Specimen of *Litus* nr. *camptopterus* (QMOR88023).

### Distributional records

We found several new species records for both the province of Quebec and the country of Canada, according to the GBIF database and the latest North American checklist (Huber et al. 2021). *Alaptus terebrans*, *Alaptus iceryae*, *Alaptus antennatus*, *Anagrus mockfordi*, *Camptoptera magna* and *Platystethynium vagatum* are newly recorded in Canada. *Camptoptera papaveris*, *Erythmelus burtik*, *Lymaenon litoralis*, as well as every *Alaptus* species identified are new records for Quebec.

### DNA barcode databases

We identified our 2,098 sequences with BLAST searches on GenBank and BOLD, which returned four specific identifications for 107 specimens: *Cosmocomoidea morrilli* (Howard) (*Gonatocerus morrilli* in BOLD; 3 individuals), *Anaphes listronoti* Huber (39 ind.), *Anagrus ustulatus* Haliday (2 ind.) and *Anagrus daanei* (73 ind.). Except for two specimens identified as *A. ustulatus* (now synonymized with *A. atomus* (Triapitsyn et al. 2020)), no species identifications matched ours. The three individuals we identified as *A. daanei* were not part of the ones identified to this species in the BOLD database. *Cosmocomoidea morrilli* determinations in BOLD were confirmed to be erroneous: our specimens were from the genus *Gonatocerus* sensu stricto. *Cosmocomoidea morrilli* has antennae overall dark but with the 5th and 6th segments distinctively white, as well as a basal brown spot on the forewings (Triapitsyn, Huber, et al. 2010). According to the photographs in BOLD, none of the specimens associated with the BIN of *C. morrilli* (BOLD:AAU9165) possess these characteristics. *Anaphes listronoti* can be distinguished in part by having multiporous plate sensillae on the second antennal segment (Huber et al. 1997), which our specimens did not have. The remaining 1,668 and 319 specimens had accurate generic and family identifications, respectively.

We added species names to 16 previously unidentified Mymaridae COI barcodes in the reference databases: *Alaptus minimus*, *A. terebrans*, *A. antennatus*, *A. iceryae*, *A. immaturus*, *A. huberi*, *Anaphes luna*, *A. gerrisophaga*, *Camptoptera magna*, *Dicopus halitus*, *Erythmelus burtik*, *Eustochus pengellyi*, *Gonatocerus rivalis*, *Lymaenon litoralis*, *Platystethynium vagatum*, *Ptilomymar magnificum*.

## Discussion

This study aimed at discovering the diversity of Mymaridae in temperate forests of Quebec using megabarcoding and integrative taxonomy. This is the first large-scale study of this dark taxon, as well as the first attempt to test the reverse workflow with the LIT criteria on Hymenoptera.

### DNA megabarcoding

DNA megabarcoding (Chua et al. 2023) was very successful, with 98% of our specimens obtaining a minibarcode. This method maintains the link between the specimen and its DNA sequence, in contrast to metabarcoding. From our personal experience, Mymaridae DNA can be difficult to amplify with universal primers commonly used for mini barcodes, such as mlCO1intF (Leray et al. 2013) and jgHCO2198 (Geller et al. 2013). The new primer combination selected in this study revealed to be more effective for mymarids and should be employed in future research aiming to sequence this taxon. Furthermore, as in Cruaud et al. (2017), we were able to obtain DNA sequences from specimens that showed no bands on the electrophoresis gel. The low cost of next generation sequencing (NGS) make it advantageous to try sequencing those amplicons. NGS also permits the detection of coamplification of non-target sequences like human contamination, endosymbiotic bacteria such as *Wolbachia*, pseudogenes and heteroplasmy (Song et al. 2008, Smith et al. 2012, Cruaud et al. 2017). We encountered these cases during the bioinformatic analyses. We were able to filter out these non-targets and keep only the target sequences, which would not have been possible with Sanger sequencing.

### Integrative species delimitation

Overall, ASAP performed better than RESL, which greatly overestimated the number of putative species of Mymaridae in our dataset. Similar overestimated results with RESL were obtained for other insect taxa (Ranasinghe et al. 2022, Pentinsaari et al. 2017, Song et al. 2018). The partition of ASAP42 was the closest to the morphologically recognized species richness of our sample. Six incongruent clusters had to be split into multiple species, of which two matched the delimitation by ASAP83. For the remaining four incongruent clusters, we adopted a conservative approach to species delimitation: we did not split species when DNA and morphology did not lead to unambiguous differentiation. Excess lumping of some species will be easier to resolve in future studies than excessive splitting (Carstens et al. 2013, Miralles and Vence 2013). We also chose not to include a tree-based delimitation method since shorter sequences can lead to errors in phylogenetic estimations and thus failure in species delimitations (Reid and Carstens 2012, Zhang et al. 2013).

A large number of haplotypes and species that are difficult to identify have been shown to negatively affect species delimitation with ASAP (Magoga et al. 2021). MOTUs with large numbers of specimens and high intraspecies genetic divergence can also have a greater probability of being incongruent with morphology (Meier et al. 2025). Our two most problematic clusters, 1.1 (*Alaptus minimus*, N=930) and 2.5 (*Anagrus* sp., N=83), contained 65 and 51 haplotypes respectively. We suggest that these two clusters contain multiple species due to their high genetic diversity, but the available morphological data do not permit their delimitation. A few species have been synonymized with *A. minimus* over the years (Triapitsyn 2017) and they might be represented in this broad genetic divergence. Having access to type specimens and their DNA would help to decide if they deserve to be re-established as valid species. Unfortunately, it is not possible to extract DNA from specimens mounted on slides. The fact that three other *Alaptus* clusters keyed to *A. minimus* suggests that morphological characters currently used to separate species might not be precise enough to reflect the diversity of this genus. *Anagrus* species are also known to be particularly difficult to determine, with high morphological polymorphism and the presence of many undescribed species (Triapitsyn 2015, Zanolli et al. 2016). These different factors could potentially explain the challenges encountered with their delimitation.

Now that it is relatively easy to obtain DNA sequences at lower cost, multilocus approaches are generally preferred to single locus methods for species delimitation (Carstens et al. 2013, Fontaneto et al. 2015). Using a second genetic marker, such as a nuclear gene, might help resolve the more complex cases like the *A. minimus* and *Anagrus* sp. clusters. It could also help resolve potential cases of heteroplasmy or hybrid introgression. The endosymbiotic bacteria *Wolbachia* can be introduced in the mitochondrial genome by hybrid introgression (Raychoudhury et al. 2009). Sequences of *Wolbachia* were found in some of our amplicons, mainly from the genus *Alaptus*. Bacterial endosymbionts have been shown to reduce the efficacy of DNA barcoding for Pteromalidae (Hymenoptera: Chalcidoidea) (Raychoudhury et al. 2009) and to lead to high genetic divergence between infected and non-infected populations of fig wasps (Chalcidoidea: Agaonidae) (Xiao et al. 2012). A multilocus approach could help decipher if this is a problem affecting

Mymaridae. Nonetheless, COI was efficient at delimiting many of the species found in our sampling and is a useful first step when working with mostly unknown taxa.

### The LIT approach

Overall, the reverse workflow with the application of the LIT criteria (Hartop et al. 2022) was successful in accelerating the discovery process of mymarid species. Pre-sorting with MOTUs was advantageous compared to the traditional approach of morphological sorting. Most of the important morphological characters need to be precisely measured to differentiate between species and this cannot be done without labor intensive slide-mounting. Since species are sexually dimorphic and most are only identifiable by their females, morphological sorting excludes a large part of the samples. Molecular sorting is thus aptly suited to mymarids.

Only the stability criterion turned out to be important in flagging PI clusters. Every PI cluster flagged exclusively with a p-distance larger than 1.5% was consistent with morphology. It is possible that this threshold was too low to indicate PI clusters for Mymaridae, since a few confirmed species reached a maximum divergence of 4-5%. High intraspecific genetic distances are not rare within Hymenoptera, with almost a third of species exceeding 3% (Zhang and Bu 2022), and some even going up to 14% (Schmidt et al. 2017). We would suggest the use of only the stability criterion in future studies within Hymenoptera. Nonetheless, five clusters were flagged solely by high divergence and resulted in just five additional specimens to validate. In total, only 9% (N=185) of the sample had to be mounted on slides for morphological evaluation.

### DNA barcode databases

As we have seen from our results, Mymaridae are largely underrepresented in barcoding databases. Only 5% of our specimens received a specific determination, of which all but two ended up being erroneous anyway; less than 1% of our sequences were correctly identified using DNA reference libraries. These results are similar to previous research, with only 1% of microhymenoptera assigned to species level (Quicke et al. 2023, Geiger et al. 2006). Errors in databases are common (Janko et al. 2024, Mioduchowska et al. 2018, Cheng et al. 2023), and taxonomic misidentifications can have negative impacts for conservation and biological control (Bortolus 2008, Bickford et al. 2007, Driesche and Hoddle 2016). Major improvements are needed if the initial goal of DNA barcoding, the identification of species, is to be useful for this family. Nonetheless, our study provides well-curated and identified novel COI barcodes for 16 described species of mymarids. This improvement will help to overcome their taxonomic neglect and make more precise biodiversity assessments of this most abundant dark taxa with approaches such as metabarcoding or eDNA.

### Mymaridae diversity

We recovered 16 mymarid genera, four of which were not previously recorded in Quebec according to the latest checklist by Huber et al. (2021). Seven of the genera known to occur in Quebec were not collected, at least in the subsample used for this study. Some genera were much more abundant than others, namely *Alaptus*, *Dicopus* and *Anagrus* (Table 2). The only other study that looked at Mymaridae diversity in forested environments in Canada found similar abundance results (Vance et al. 2007), with *Alaptus* largely overrepresented. Unfortunately, but for reasons discussed here, they did not identify their specimens below genus level.

We delimited a total of 55 species, almost doubling the richness found in Quebec from the sampled genera (Table 2) (Huber et al. 2021). We believe that a few of our challenging clusters ultimately consist of multiple cryptic species that we were not able to delimit with the data at hand. Consequently, the true richness is likely higher than what we have estimated. Since 13 clusters could not be confidently identified to named species, they might represent species new to science. The 18 clusters that were morphologically close to described species might also contain some new species. However, several of those were represented by one or a few individuals so more specimens would need to be evaluated before committing to a formal description. Comparisons with type specimens would also be required to confirm their validity.

Twenty-one of the species identified have a known geographic distribution outside of Canada, with some outside the North American continent (Huber et al. 2021). With the fact that Mymaridae attack host eggs that are mostly concealed in plants or soil, they can be easily introduced in other habitats with trade (Triapitsyn 2015). Most species probably have a widespread distribution. It is thus possible to find species only described from other countries.

Ultimately, our study contributes to the knowledge of Mymaridae from Quebec for the first time. Our results confirm the effectiveness of the LIT method for taxa other than Diptera. Besides the clear advantage of LIT in speeding up the taxonomic process, it also establishes criteria that are applied in a systematic and reproducible manner. Results also show that an integrative taxonomic approach is essential for dark taxa like microhymenoptera, because neither morphology nor DNA alone would have recovered this diversity. Further research is still needed to resolve conflicts that occurred in our study, with the evaluation of more specimens and possibly of a third type of data, such as nuclear DNA. Still, the amount of data we have contributed to reference databases will help future biodiversity inventory and conservation of parasitoid wasps.

## Author contributions

CH: Conceptualization; Data curation; Formal analysis; Investigation; Methodology; Visualization; Writing – original draft. CF: Conceptualization; Funding acquisition; Methodology; Resources; Supervision; Writing – review & editing

## Acknowledgements

We thank Dr. John Huber for his time, advice and access to specimens from the CNC. We also thank the volunteers at the Ouellet-Robert Entomological Collection, Martin Lefrançois, Marianne Goulet and Malek Kalboussi (Université de Montréal) for their help with the project. We acknowledge the support of the Natural Sciences and Engineering Research Council of Canada (NSERC) for a Canada Graduate Scholarship-Master’s and a *Fonds de recherche du Québec – Nature et Technologies* scholarship (https://doi.org/10.69777/329893) to C. Hébert. Research funding was provided by NSERC [funding reference number RGPIN-2024-05502].

## Data Availability

Machine-readable data for 2098 natural history specimens are available in Hébert and Favret (2025). DNA sequences taken from those same specimens are available on BOLD (dx.doi.org/10.5883/DS-MYMAQC) and GenBank (accession numbers PX064719-PX066816). The benefits of this research include the sharing of our data on public repositories as outlined above.

